# The accuracy of LD Score regression as an estimator of confounding and genetic correlations in genome-wide association studies

**DOI:** 10.1101/234815

**Authors:** James J. Lee, Matt McGue, William G. Iacono, Carson C. Chow

## Abstract

In order to infer that a single-nucleotide polymorphism (SNP) either affects a phenotype or is linkage disequilibrium with a causal site, we must have some assurance that any SNP-phenotype correlation is not the result of confounding with environmental variables that also affect the trait. In this work we study the properties of LD Score regression, a recently developed method for using summary statistics from genome-wide association studies (GWAS) to ensure that confounding does not inflate the number of false positives. We do not treat the effects of genetic variation as a random variable and thus are able to obtain results about the unbiasedness of this method. We demonstrate that LD Score regression can produce estimates of confounding at null SNPs that are unbiased or conservative under fairly general conditions. This robustness holds in the case of the parent genotype affecting the offspring phenotype through some environmental mechanism, despite the resulting correlation over SNPs between LD Scores and the degree of confounding. Additionally, we demonstrate that LD Score regression can produce reasonably robust estimates of the genetic correlation, even when its estimates of the genetic covariance and the two univariate heritabilities are substantially biased.

## 1 Introduction

The goal of genome-wide association studies (GWAS) is to find regions in the genome where variation affects a phenotype. However, this must be accomplished from observed correlations, and inferring causation from correlation is a famously perilous endeavor (Freedman, 1999; Pearl, 2009). The GWAS field has been fortunate in that it offers a variety of methods to check whether confounding effects have produced spurious correlations between genetic and phenotypic variation. These methods have led to a strong consensus that confounding has a minimal impact on GWAS results (Goldstein, 2011; Visscher, Brown, McCarthy, & Yang, 2012; Lee, 2012; Lee, Vattikuti, & Chow, 2016).

One of the newer methods used to check the causal status of GWAS associations is known as LD Score regression (Bulik-Sullivan et al., 2015a), which can be applied to summary statistics assembled from the contributions of different research groups and thus does not require access to individual-level data. This technique relies on the simple linear regression of assayed single-nucleotide polymorphism (SNP) *j*’s association chi-square statistic on

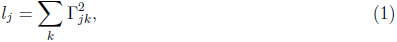

the sum over all SNPs of each SNP’s squared correlation with the focal SNP *j*. This latter quantity is called SNP *j*’s “LD Score.” Empirically, the regression curve relating chi-square statistics to LD Scores is always very close to an upwardly sloping straight line. This result is explicable because a SNP tagging more of its neighbors—and, thus, having a higher LD Score—is more likely to tag one or more causal sites affecting the phenotype. The lowest possible LD Score of a SNP is one, which is obtained when a SNP is in perfect linkage equilibrium (LE) with all other SNPs. A hypothetical SNP with an LD Score of zero fails to tag the causal effect of any SNP in the genome—including whatever effect the SNP itself may have. Therefore, if the intercept of LD Score regression departs upward from unity (the theoretical expectation of the chi-square distribution with one degree of freedom), then intuitively the departure must be due to confounding, poor quality control, overlapping samples in the meta-analysis, or other artifacts. This simple and insightful method of estimating the average chi-square statistic of truly null SNPs (or at least a certain subset of such SNPs) should in most cases lead to a much better global correction of the association statistics than the overly conservative genomic control (Devlin & Roeder, 1999).

The slope obtained from LD Score regression could in principle also provide an estimate of the trait’s heritability—the fraction of the phenotypic variance ascribable to genetic differences in the population. The developers urge caution when putting the method to this use, particularly if the heritability is not partitioned in the same analysis (https://nealelab.github.io/UKBB_ldsc/h2_univar_browser.html, accessed May 17, 2018), and we will give some reasons why LD Score regression may not accurately estimate heritability below.

Another use of LD Score regression is the estimation of genetic correlations (Bulik-Sullivan et al., 2015b). The dependent variable in this case is not the chi-square statistic from the GWAS of a single trait but rather the product of two *Z* statistics, each taken from a GWAS of a distinct trait. In principle, this use offers a means of determining whether a trait-trait correlation (as opposed to a SNP-trait correlation) is attributable to the presence of confounders affecting both traits. If the genetic correlation is statistically and quantitatively significant, then we can be sure that the total phenotypic correlation is not attributable solely to confounders that are entirely environmental in nature. Many interesting relationships have been confirmed or discovered by bivariate LD Score regression, including a high genetic correlation (∼0.70) between years of education and age at first childbirth (Barban et al., 2016) and a moderate one (∼0.35) between years of education and intracranial volume (Okbay et al., 2016).

In the classical era of quantitative genetics, genetic correlations were most commonly estimated with twin data. Rather large samples of twinships are required for precise estimates with this design, and in some cases the estimates are not as robust against modeling assumptions as estimates of univariate heritabilities (Beauchamp, Cesarini, Johannesson, Lindqvist, & Apicella, 2011). For these reasons a welcome development in quantitative genetics has been the advent of GWAS, which can now reach sample sizes in the hundreds of thousands. The appearance of robustness offered by GWAS can be illusory, however, if estimates of genetic correlations are themselves subject to confounding. One can devise estimators of the genetic correlation that might be biased by environmental confounders that affect both phenotypes and happen to be correlated with genetic variation (Palla & Dudbridge, 2015; Okbay et al., 2016). An attractive feature of LD Score regression in this respect is that its control of confounding extends not just to the evidence of association at individual SNPs but also to its genome-wide estimates of genetic correlations. This is important because, again, it is precisely the issue of a phenotypic correlation’s underlying causal nature that can call for an accurate estimate of the genetic correlation.

As appealing as the intuition behind LD Score regression may be, the mathematical justifications of this method given so far in the literature raise questions because of their assumption that the effects of genetic variants can be treated as a random variable. This assumption is a useful convenience for computations, but it is not biological. The effects of genetic polymorphisms should be invariant; it is genotypes and phenotypic residuals that vary between individuals (Lee & Chow, 2014; de los Campos, Sorensen, & Gianola, 2015). The assumption also precludes a quantitative treatment of the method’s accuracy. Here we refrain from this assumption of random genetic effects and instead treat the effects as a vector of arbitrary fixed constants. Hence we are able to obtain precise expressions of the quantities estimated by LD Score regression, which can be compared with the quantities of actual interest to determine when they coincide. Here is a preview of our results:

1. If the effects of the standardized genotypes at SNP *j* and its correlated neighbors is not related to SNP *j*’s LD Score, then the slope of LD Score regression provides an unbiased estimate of heritability. For both biological and evolutionary reasons, however, genetic effects are typically smaller near SNPs with higher LD Scores (Gazal et al., 2017). LD Score regression may therefore not be a reliable way to estimate the heritability of a trait (or, by extension, the genetic covariance between two traits).
2. The intercept of LD Score regression reflects a useful measure of confounding in the GWAS even in an important case of a relationship between LD Scores and the correlations of SNPs with environmental confounders. This is perhaps the most novel and important conclusion of our analysis. The developers of LD Score regression warn that in the general case of such a relationship the intercept will not accurately estimate the contribution of confounding to the GWAS statistics (Bulik-Sullivan et al., 2015a). One reason for such a relationship, however, is that the genotypes of the parents have an effect on the phenotype of the offspring that is not mediated by the offspring’s own genotype. A prominent example of this phenomenon is parents with a genetic disposition to obtain more education creating an environment for their offspring that also promotes educational attainment (Sacerdote, 2007; Kong et al., 2018; Lee et al., in press). In this special but important case, the intercept of LD Score regression can still be used to correct the association statistics of null SNPs so that their average chi-square statistic is in line with the null hypothesis of no causality.
3. LD Score regression provides an accurate estimate of the genetic correlation between two traits, even if neither trait’s heritability is well estimated.

## 2 Materials and methods

In the Supplementary Note, we present a mathematical analysis of LD Score regression’s important properties that does not treat the average effects of gene substitution as random variables. To confirm our mathematical results, we conducted simulations using the Minnesota Center for Twin and Family Research (MCTFR) genetic data (Miller et al., 2012). The MCTFR cohort consists of 8,405 participants, clustered in families, each typically consisting of a father, mother, and two twin offspring. All cohort members were genotyped at 527,829 SNPs with the Illumina Human660W-Quad array. The other genotypes of the European-ancestry cohort members were subsequently imputed (1000 Genomes phase 1), producing calls at more than 8 million SNPs with a relatively high minor allele frequency (MAF). In the imputation step, data was obtained from only one member of each monozygotic (MZ) twinship, which led to a total sample size of roughly 6,700. There is a large degree of overlap between these imputed SNPs and those used in the calculation of LD Scores by Bulik-Sullivan et al. (2015a).

Our first set of simulations was intended to study the relationship between the LD Scores of causal SNPs and estimates of heritability. To minimize computational burden, we calculated our own MCTFR-specific LD Scores, using the Illumina genotyping data from the ∼4,000 parents. We limited the summation in Equation (1) to SNPs within the recommended 1-cM window of SNP *j*. We collected these LD Scores into one file and examined the quantiles of their distribution. We called the LD Scores below the 25th percentile (4.744) *very low*, those between the 25th percentile and the median (7.154) *low*, those between the median and the 75th percentile (10.47) *high*, and those above the 75th percentile *very high*. Any given simulation condition used a sample of 5,000 causal SNPs from either just one of these categories or at random from all ∼500,000 genotyped SNPs, assigning them a normal distribution of standardized effects (Fisher, 1941; Lee & Chow, 2013) such that the total heritability equaled 0.50. We used PLINK 1.9 (Chang et al., 2015) to carry out a GWAS of the simulated phenotype. We then applied LD Score regression (downloaded September 2016 from https://github.com/bulik/ldsc) to the GWAS statistics to estimate the intercept and the heritability. A hundred replicates were conducted of each condition (*very low*, *low*, *random*, *high*, *very high*), each time keeping the same vector of average effects sampled for that condition but assigning the ∼4,000 subjects different non-genetic residuals. In this set of simulations only the MCTFR white parents were used as subjects.

We retained this simulation framework to study the accuracy with which bivariate LD Score regression estimates genetic correlations. Here we did not sample causal SNPs from just one quartile of LD Score, because in certain conditions this would preclude any nonzero genetic correlation. To simulate a genetic architecture tending to produce unbiased estimates of genetic covariance and heritability, we assigned all genotyped SNPs in MCTFR an average effect drawn from a normal distribution. To simulate a genetic architecture where lower-LD SNPs have larger effects, we multiplied the effects of all SNPs with the *low* annotation by two and then rescaled the vector of effects so that it satisfied the target heritability (0.8). Conversely, to simulate a genetic architecture where higher-LD SNPs have larger effects, we multiplied the effects of all SNPs with the *high* annotation by two and then rescaled. Draws from the bivariate normal distribution were used to induce the desired genetic correlation between the two traits. In scenarios where the two traits were related to LD in opposite ways, the correlation parameter of the bivariate normal distribution was fixed to be higher than the target genetic correlation so that the latter would end up being the correlation between the two vectors of average effects after the multiplications of effects at disjoint SNPs. In all conditions we fixed the total heritability to an unrealistically high 0.8, because preliminary runs with a heritability of 0.5 sometimes led to the software returning an error rather than taking the square root of a negative heritability estimate. Four thousand subjects is a small sample by the standards of LD Score regression (http://ldsc.broadinstitute.org/upload_file).

Our final set of simulations was intended to study whether the intercept continues to be an effective means of controlling the Type 1 error rate with respect to the null hypothesis that the SNP is neither causal nor in LD with a causal site, in a certain case of LD-dependent confounding. Here the MCTFR white offspring were used as the subjects in the simulated GWAS. To compensate for the resulting reduction in sample size, we both increased the number of replicates in each condition to 200 and used the precomputed whole-genome LD Scores available from https://data.broadinstitute.org/alkesgroup/LDSCORE. The latter step ensured a greater number of observations (SNPs) in the regression of chi-square statistics on LD Scores. We made all imputed SNPs on odd chromosomes causal and all imputed SNPs on even chromosomes non-causal; the SNPs on even chromosomes were thus rendered suitable for examining the Type 1 error, since they were all guaranteed to be null. In the conditions intended to simulate confounding that increases with LD Score, we took half the average breeding (additive genetic) value of each offspring’s parents and added it to the part of the offspring’s non-genetic residual that was independent of breeding value. (This latter part always had a variance of 0.5.) That is, using the same genetic architecture determining the “true polygenic scores” of the offspring, we calculated the true polygenic scores of the two parents in a family and treated the average as an environmental variable affecting the offspring phenotype with a path coefficient of 0.5. A path diagram representing this causal system is presented in Figure 1.

**Figure 1:**
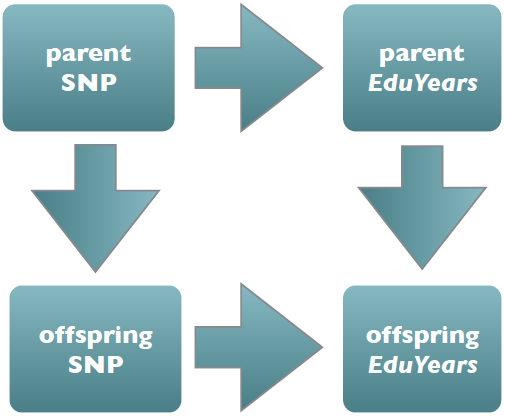
A causal graph (path diagram) displaying a mechanism where the extent of confounding increases with LD Score. Evidence for this mechanism has been uncovered in recent studies (Kong et al., 2018; Lee et al., in press). See the Supplementary Note for a more detailed explication.

It is desirable to combine this special form of confounding with a more conventional form envisaged by GWAS investigators. An ideal way to simulate population stratification might be to use two cohorts sampled from opposite ends of Europe (or analyzed with different genotyping/imputation pipelines) and to give each cohort a different mean residual. We refrained from using principal components as a proxy for such structure because a subject’s projection on a principal component is simply a linear combination of genotypes (Price et al., 2006). If the projection is then used as a basis for how to perturb the phenotype, it becomes very difficult to say how the simulated process is any different from a true causal effect of genotype on phenotype. Because we lack any way of discerning structure within the MCTFR whites independently of principal component analysis, we were forced to simulate bias through another means that happens to be highly convenient in MCTFR—the inclusion of close relatives in the sample. In these simulation conditions, we augmented the sample with 694 individuals, each of whom is a dizygotic (DZ) twin of an original sample member. This raised the offspring sample size to 2,701. (At the outset we chose a twin at random from each DZ twinship to create a sample of unrelated individuals. In the conditions incorporating relatedness, we thus brought back the twin who was initially excluded.) Relatedness induces a spurious inflation of the GWAS chi-square statistics because the effective sample size is not as large as it seems. Note that when relatedness is combined with an effect of parent genotype on offspring phenotype (Figure 1), relatedness additionally becomes a kind of population stratification. Each family is its own population, represented by as many two members in the sample, and even null SNPs become associated with the phenotype because they are indicative of parentage and thus of a key environmental factor affecting the phenotype.

As a robustness check, we reran the final set of simulations with the heritability set to zero. The average of the parental phenotypic values was used instead of their genetic values to perturb the offspring phenotypes, since a zero heritability by definition implies that no one has a genetic value. Note that a combination of relatedness and an effect of parent phenotype is still a form of population stratification even in this case of a non-heritable trait.

## 3 Results

### 3.1 The slope of univariate LD Score regression as an estimator of heritability

In the Supplementary Note, we show that the slope of LD Score regression provides an unbiased estimate of the heritability if a SNP’s LD Score is unrelated to the per-SNP heritability of the SNP itself and its LD partners. The requirement of this null correlation for an unbiased estimate of heritability is stringent. Regressing chi-square statistics on LD Scores to estimate the heritability depends on a constant average per-SNP heritability regardless of LD. If average per-SNP heritability declines in higher-LD regions, say, then the estimated heritability must fall short of the true heritability. This sensitivity to LD is a feature shared with the heritability-estimation method GREML (Speed, Hemani, Johnson, & Balding, 2012; Lee & Chow, 2014; Yang et al., 2015; Chen, 2016).

A negative correlation between LD and heritability tagged per SNP may well be the rule (Gazal et al., 2017), for at least two reasons. First, if the region surrounding the focal SNP is under evolutionary constraint, then mutations occurring at nearby sites will typically be eliminated by selection and thereby fail to become present-day SNPs contributing to the focal SNP’s LD Score. Second, the higher recombination rate in functionally important regions, such as those that are DNase I hypersensitive, leads to a more rapid attenuation of LD between the focal SNP and the neighboring polymorphisms that do manage to persist over evolutionary time. In this case of SNPs with higher LD Scores tagging less heritability, the slope of LD Score regression leads to an underestimation of the true heritability.

**Table 1:**
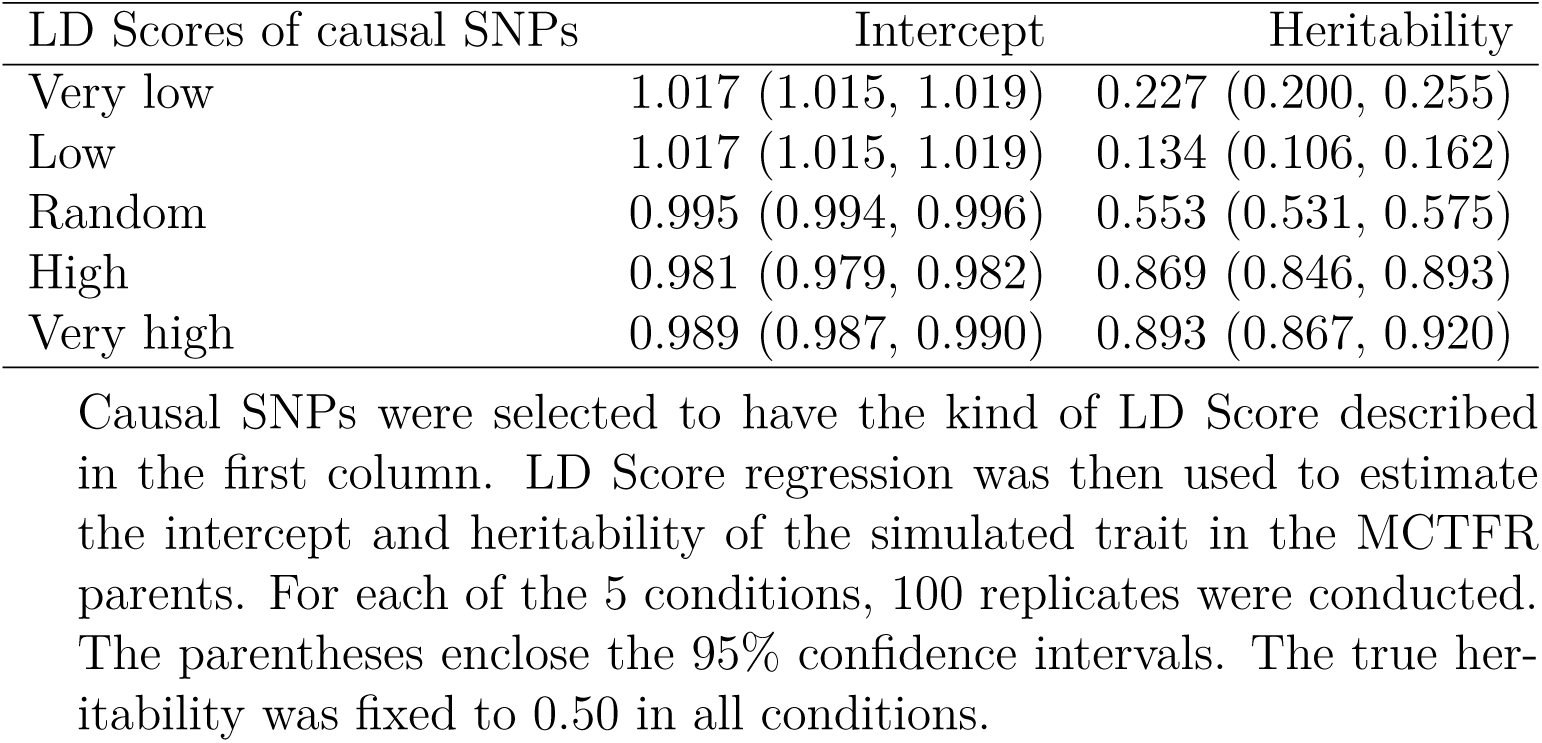
Properties of univariate LD Score regression in simulations based on the MCTFR genetic data

| LD Scores of causal SNPs | Intercept | Heritability |
| --- | --- | --- |
| Very low | 1.017 (1.015, 1.019) | 0.227 (0.200, 0.255) |
| Low | 1.017 (1.015, 1.019) | 0.134 (0.106, 0.162) |
| Random | 0.995 (0.994, 0.996) | 0.553 (0.531, 0.575) |
| High | 0.981 (0.979, 0.982) | 0.869 (0.846, 0.893) |
| Very high | 0.989 (0.987, 0.990) | 0.893 (0.867, 0.920) |

Causal SNPs were selected to have the kind of LD Score described in the first column. LD Score regression was then used to estimate the intercept and heritability of the simulated trait in the MCTFR parents. For each of the 5 conditions, 100 replicates were conducted. The parentheses enclose the 95% confidence intervals. The true heritability was fixed to 0.50 in all conditions.

We conducted a set of simulations to test these theoretical deductions. We chose the causal SNPs either randomly or on the basis of their LD Scores and studied the impact of this choice on estimates of the heritability. The results are displayed in Table 1. A random selection of causal SNPs led to an average estimate of heritability (0.553) reasonably close to the true *in silico* heritability (0.50). The relationship between LD dependence and heritability estimate appears to be non-monotonic, and we will shortly discuss possible reasons for this. Nevertheless there is an overwhelmingly evident trend for the heritability estimates to be too low when the causal SNPs all have below-median LD Scores (and conversely too large when the causal SNPs all have above-median LD Scores), in accordance with our theory.

Our simulations testing the accuracy of bivariate LD Score regression as an estimator of genetic correlations produced as byproducts estimates of the two univariate heritabilities in each run, and Table S1 presents the results. Surprisingly, estimated heritabilities varied substantially, beyond what is expected as a result of sampling error, even under the same values of the simulation parameters. For example, even though the average effects were drawn from the normal distribution and then rescaled in the same way, the estimated heritabilities of traits with a low-LD bias ranged from 0.28 to 0.62. It may be that even the small fluctuations in the relationship between LD and effect size induced by our scheme for generating the genetic architecture can have substantial effects on the heritability estimate returned by LD Score regression. (Recall that in a given condition we did not redraw the average effects of the SNPs for a new replicate. We only redrew the non-genetic residuals of the individuals.) It has also been suggested to us that the reason for the instability may be the small size of the simulation sample (∼4,000), which falls below the recommendation of the developers. Nevertheless we can see that the overall results bear out our theoretical arguments. The average of the estimates over all *unbiased heritability* conditions is 0.803, extremely close to the true *in silico* value of 0.8. The average of the estimates over all conditions intended to induce an upward bias is 0.842. The average of the estimates over all conditions intended to induce a downward bias is 0.451, suggesting an asymmetrically greater sensitivity to downward rather than upward bias.

In summary, even though the simulations whose results are presented in Tables 1 and S1 assigned effects to SNPs in markedly different ways, they jointly affirmed that a dependence of per-SNP heritability on LD Score leads to inaccurate estimates of overall heritability.

### 3.2 The intercept of univariate LD Score regression as an estimator of confounding

A far more important use of LD Score regression is the estimation and correction of confounding (or any other bias that can inflate the association statistics, such as overestimation of the effective sample size as a result of close relatives in the sample). If the intercept of LD Score regression is truly equal to the average chi-square statistic of SNPs that neither affect the phenotype nor tag any causal sites, then dividing all of the GWAS chi-square statistics by the intercept should restore the average chi-square statistic of these null SNPs to the theoretically proper value of unity and bring the Type 1 error rate close to the targeted level. We now examine the extent to which this use of the method is valid.

We first suppose that the magnitude of a SNP’s correlation with environmental factors affecting the phenotype is independent of its LD Score. Such independence implies that the conditional average increase in the chi-square statistic due to confounding at each possible LD Score does not in fact vary as a function of LD Score, and thus the entire regression line is elevated by a uniform amount. The intercept is expected to be very close to unity in the absence of confounding (Table 1; Figures 2 and S1), and therefore the amount by which the regression line is moved upward can be determined from the departure of the intercept from unity. Furthermore, suppose that null SNPs do not differ from non-null SNPs in the average extent of confounding—which is extremely likely if LD Scores are indeed independent of confounding. After all, SNPs differing in LD Score also differ in their probability of being null (the probability increasing as the LD Score declines), and it is hard to see how the same spurious increase in the chi-square statistic can be maintained as the LD Score varies (and hence as the mixture of null and non-null proportions varies)—unless null SNPs do not in fact differ from non-null SNPs in the extent of their confounding with environmental factors. It follows that null SNPs have an average chi-square statistic equal to the intercept, and division of all chi-square statistics by the intercept will bring their average back to the required value of unity. This conclusion was also reached by Bulik-Sullivan et al. (2015a).

We will now show that division by the intercept can still be viable means of correcting confounding in some situations where LD Scores and SNP-environment correlations are related.

Suppose that the spurious increase in the chi-square statistic depends linearly on LD Score. This is extremely likely if the regression of the total chi-square statistic on LD Score is linear, since it would be quite a coincidence if the superposition of terms with markedly nonlinear relationships with LD Score produced a closely linear relationship. SNPs with an LD Score of one can tag at most one causal SNP, SNPs with an LD Score of two can tag more than one, and so on. The linear extrapolation to an LD Score of zero represented by the intercept is thus very close to the confounding-induced inflation of the chi-square statistics at SNPs that are null by virtue of tagging very few SNPs. If the trait is not sufficiently polygenic, then there will be many SNPs with moderate or large LD Scores that happen to be null. Then the intercept reflects the average confounding-induced inflation at only a certain subset of null SNPs, and one might worry that the chi-square statistics of null SNPs outside of this subset are not properly corrected.

An important case of such dependence is a direct (non-genetic) effect of parent on offspring phenotype, such as when highly educated parents can help their offspring (even if adopted) become highly educated in turn (Sacerdote, 2007). Figure 1 depicts this situation. The causal effect of the offspring’s own genotype on phenotype will be accurately estimated in within-family studies (Laird & Lange, 2006; Lee & Chow, 2013), but the within-family estimate will fall short of the estimate obtained from GWAS of unrelated individuals because the latter also reflects the confounding influence of the parent genotype. Although we have depicted the parent years of education as the mediator of the parent genotype’s causal influence (above and beyond its influence through the offspring genotype), the mediator does not necessarily have to be the same phenotype studied in the GWAS of the offspring. When the offspring phenotype is years of education, parent characteristics acting as mediators might also include intelligence, income, and other determinants of social status (Clark, 2014). Our results below are applicable whenever there is a high genetic correlation between the offspring phenotype and the mediating parent characteristic (Marioni et al., 2014; Hill et al., 2016).

In this case the spurious increase in a SNP’s GWAS coefficient is equal to the square of the true genetic coefficient up to a constant factor, plus whatever part of the spurious increase does not depend on the true coefficient (Lee, 2012). Critically this affine dependence means that null SNPs, regardless of their LD Scores, will have an average chi-square statistic equal to the intercept so long as the confounding that is independent of the true genetic coefficient does not depend on LD Score. Division of all chi-square statistics by the intercept again leads to the subset of statistics corresponding to null SNPs having the required average of unity. Note that the inability to factor out the contribution of confounding to the chi-square statistics of non-null SNPs in these cases simply leaves us with more or less statistical power to detect such SNPs without affecting the Type 1 error rate.

We provide more justification of this argument in the Supplementary Note. Here, we use simulations based on our MCTFR genetic data to provide further support. Briefly, we conducted GWAS of a simulated phenotype potentially affected by the genotypes of the parents (Figure 1) and applied LD Score regression to the resulting summary statistics. Figure 2 displays the results. One consequence of augmenting the sample with DZ twins of the original sample members is that the estimated heritability increased even in the condition with no confounding. This may be the result of SNPs with higher LD Scores having even more LD partners as a result of the “bottleneck” imposed by recent common ancestry, an effect analogous to the increase in heritability estimates obtained with GREML in samples with relatedness (Vattikuti, Guo, & Chow, 2012; Zaitlen et al., 2013). This should not affect our conclusions. What is important is that in both the *unrelated* and *DZ twin* conditions, the heritability estimate increased upon allowing the parent genotype to have an environmentally mediated effect on the offspring phenotype (*P* < 0.002). This form of confounding thus contributes more to the chi-square statistics of the SNPs with the largest LD Scores.

**Figure 2:**
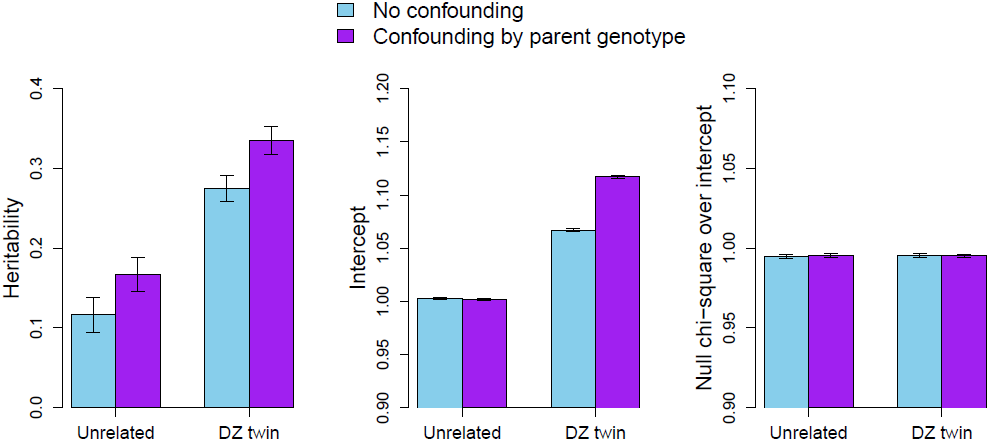
The performance of the LD Score regression intercept in simulations based on the MCTFR genetic data. Each bar and its error correspond to the average of the quantity over 200 replicates and 95% confidence interval. *Unrelated* is a sample of 2,007 nominally unrelated offspring; *DZ twin* augments that sample with 694 individuals, each of whom is a dizygotic twin of an original sample member. In the *confounding by parent genotype* condition, half the average of the two parental breeding (genetic) values was added to the offspring’s phenotype. The left panel displays the estimate of heritability based on the regression slope, the center panel displays the intercept, and the right panel displays the mean chi-square statistics of SNPs on even chromosomes (simulated to be non-causal) divided by the intercept.

In the *unrelated* conditions, the intercept remained close to unity even when there was confounding by parent genotype. This is consistent with our argument that the intercept is unaffected by this type of confounding despite dependence on LD Score. In the *DZ twin* conditions, the intercept increased as a result of the relatedness between sample members. It further increased upon the addition of confounding by parent genotype. This is consistent with relatedness becoming a type of population stratification when combined with an effect of parent genotype on offspring phenotype; each family is its own population, and even null SNPs become associated with the phenotype because they are indicative of parentage and thus of a key environmental variable affecting the phenotype. Crucially, however, in both the *no confounding* and *confounding by parent genotype* conditions at this level of the *DZ twin* factor, the intercept increased by almost exactly the right amount to offset the inflation of the chi-square statistic at null SNPs. In the rightmost panel of Figure 2, we can see that the chi-square statistics of SNPs on even chromosomes (simulated to be non-causal) became very close to unity upon division by the intercept, irrespective of relatedness and confounding by parent genotype. Figure S1 displays the simulation results in the case of a trait with zero heritability; here again, in all conditions, the chi-square statistics of SNPs on even chromosomes were very close to unity upon division by the intercept.

We now turn to an important caveat. Our argument concerning the robustness of the intercept depends on the linearity of the LD Score regression. It is certainly possible to create gross violations of linearity in simulations (Bulik-Sullivan et al., 2015a, Supplementary Figure 7). For example, if we depopulate high-LD regions of causal SNPs, then the regression curve can be non-monotonic, rising at first and then declining as the LD Score increases. In this case the slope of LD Score regression can be negative and the intercept greater than unity even in the absence of confounding. For this reason it is a salutary practice to inspect the binned scatterplot for any evidence of substantial nonlinearity, although unfortunately such a plot may not be informative if the sample size is small.

A mild degree of nonlinearity might have some effect on the intercept if the SNPs with largest LD Scores deviate from the linear trend extrapolated from the SNPs with the smallest LD Scores. For this reason it is fortunate that in practice LD Score regression is a weighted regression where the SNPs with the smallest LD Scores receive the largest weights. The purpose of this weighting is to address heteroskedasticity and non-independence; if the regression curve is perfectly linear, then the effect of this weighting is to improve the standard errors. If the curve is nonlinear, then an additional effect is to bring the entire regression line closer to the linear extrapolation from the SNPs with the smallest LD Scores and the intercept thereby closer to the average chi-square statistic of truly null SNPs.

A subtler kind of nonlinearity might occur if the slope provides a biased estimate of the heritability. Equation (16) of the Supplementary Note is an explicit expression for the average chi-square statistic of SNPs with an LD Score of *l*_*j*_, and setting *l*_*j*_ = 0 leads to the entire expression equaling unity. But this theoretical expression for the intercept requires the vanishing of several terms, and it may be that these terms do not diminish at the right rate for a linear regression curve to fit both the theoretical intercept and the empirical chi-square statistics. This kind of nonlinearity might have been responsible in our first set of simulations for the small but significant deviations of the intercept from unity (Table 1), which roughly tracked how the estimate of heritability deviated from the true value. In practice, however, this type of bias seems to lead consistently to an upward deviation of the empirical intercept and thus an overly conservative correction of the GWAS statistics (Loh, Kichaev, Gazal, Schoech, & Price, 2018; Yengo, Yang, & Visscher, 2018). This upward bias is in line with the biological and evolutionary reasons to expect the slope to provide an underestimate of the heritability.

Our conclusion regarding the robustness of LD Score regression as a safeguard against confounding is a novel result of our analysis. Bulik-Sullivan et al. (2015a) went to some lengths to show that LD Scores are uncorrelated with *F*_*ST*_ (a measure of population differentiation in allele frequencies) at various geographical scales within Europe. This is very convincing evidence in support of the assumption that confounding is uncorrelated with LD Scores—at least when the confounding takes the form of population stratification usually contemplated by GWAS researchers, the sampling of the individuals in the study from geographically distinct subpopulations differing in both allele frequencies and exposure to environmental factors. But we have found that even if confounding is correlated with LD Scores, the intercept of LD Score regression can still be used to ensure that null SNPs have an average chi-square statistic of no greater than unity in some important cases, including an environmentally mediated effect of the parent genotype.

With all of these considerations in mind, we turn to the recent work of de Vlaming, Johannesson, Magnusson, Ikram, and Visscher (2017a). These authors found that a very large degree of population stratification in their simulations leads to an intercept falling short of the magnitude required to restore the Type 1 error rate and also an overestimate of the heritability. Whatever the problem may be, evidence for it can be seen in their binned scatterplot of chi-square statistics and LD Scores (an average over many replicates), which shows a nonlinearity in the leftmost simulated data points that we have never observed in real empirical data. It is also worth noting that the problems in these simulations only arise when population stratification is quite extreme, leading to an intercept greater than 1.5 with rather small sample sizes. In this regime, the small multivariate fourth cumulant approximation may no longer be valid, although we think this is unlikely to be the explanation of the simulation results. In any event this example shows the importance of inspecting the binned scatterplot if this has stabilized and being cautious when the intercept is large enough to indicate substantial undiagnosed problems.

### 3.3 Bivariate LD Score regression as an estimator of genetic correlations

We now consider LD Score regression as an estimator of the genetic correlation between two traits. Previous studies have found the output of bivariate LD Score regression to be consistently close to what is produced by wholly different methods (Bulik-Sullivan et al., 2015b; Okbay et al., 2016; Shi, Mancuso, Spendlove, & Pasaniuc, 2017), and our goal now is to explain this robustness.

In bivariate LD Score regression, the slope is naively expected to be proportional to the genetic covariance. In the absence of confounding and sample overlap, the intercept is zero since the expected product of two independent and null-distributed *Z* statistics is zero. Any upward departure of the intercept from zero in this case is indicative of confounders affecting both traits, just as an upward departure from unity is analogously indicative of confounders affecting the focal trait in the univariate case. Herein lies the power of bivariate LD Score regression as a method; its estimate of the genetic correlation relies on the respective slopes of three regressions on LD Scores, the dependent variables being the product of *Z* statistics, the chi-square statistics of trait 1, and the chi-square statistics of trait 2. As a result any influence of confounders affecting one or both traits is minimized. Because of this key property, it is important to demonstrate that the estimate of the genetic correlation is reasonably accurate.

The Supplementary Note contains our mathematical treatment of this problem, including our demonstration that the estimate returned by bivariate LD Score regression is unaffected by a direct effect of parent genotype on offspring phenotype. Here we present the results of our simulations (Figure 3). Recall that the heritability estimates produced by this set of simulations varied substantially even for the same values of the governing parameters (Table S1). Such variability may affect estimates of genetic correlations as well. We can see in Figure 3, however, that the genetic correlation appears to be more robust.

**Figure 3:**
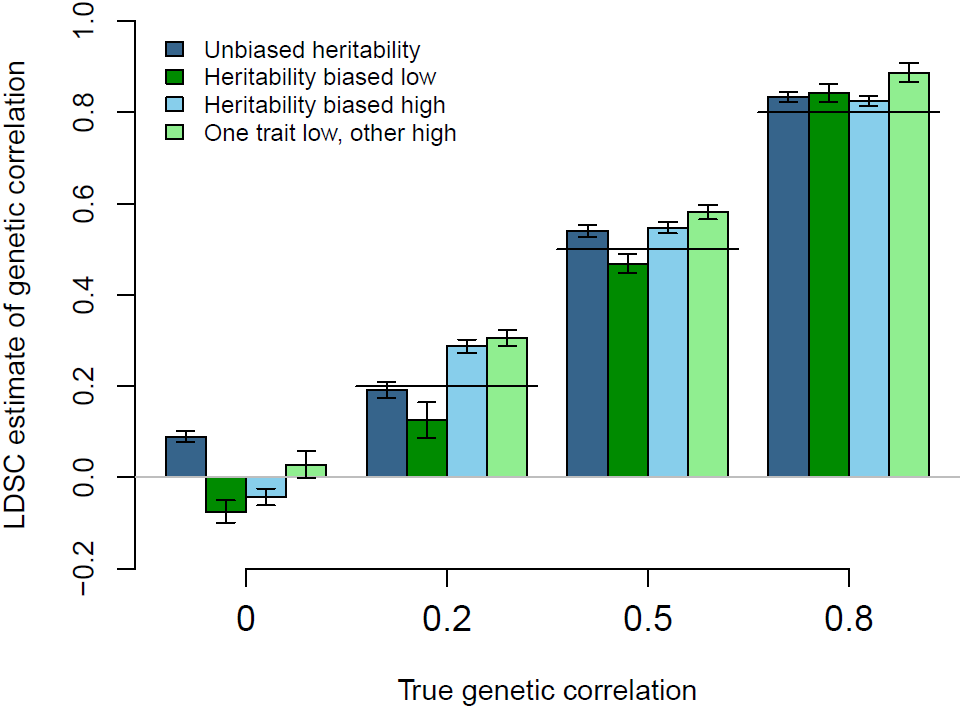
Bivariate LD Score regression was used to estimate the genetic correlation between the simulated traits in the MCTFR parents. For each condition, 100 replicates were conducted. The error bars enclose the 95% confidence intervals. The horizontal line through a given group of bars is placed at the height of the true genetic correlation.

For the most part, the simulation results affirm that bivariate LD Score regression is a robust estimator of the genetic correlation, even when estimates of the heritabilities are tremendously biased (Table S1). The largest discrepancies from the true genetic correlation, approaching 0.10 in magnitude, occurred when the correlation was fixed to the low values of zero and 0.2. Our analysis in the Supplementary Note shows that the bias in genetic correlation is expected to be larger for smaller genetic correlation, and indeed such large discrepancies did not occur once the true genetic correlation was increased to 0.5 and 0.8 (Figure 3).

## 4 Discussion

The regression of GWAS association statistics on LD Scores partitions the statistics into a part that covaries with LD Scores (the slope) and a part that does not (the intercept). Polygenic causal signal contributes to the first part by necessity, whereas confounding and other biases spuriously inflating the statistics need not make any such contribution. This insight lies at the heart of LD Score regression.

It had been presumed that in order for division by the intercept to restore the average chi-square statistic of null SNPs to the theoretically prescribed value of unity, LD Scores must be uncorrelated over SNPs with the extent of confounding with environmental influences on the phenotype. In the framework of Bulik-Sullivan et al. (2015a), this is equivalent to the absence of a correlation between LD Scores and the *F*_*ST*_ characterizing the two subpopulations. There may be such a correlation, however, in certain cases such as when the phenotype of the parent affects the phenotype of the offspring through some environmental mechanism. Remarkably we found that LD Score regression remains a robust means of correcting the association statistics, for in such a case the deviation of the intercept from unity reflects the degree of confounding at just those SNPs that are neither causal themselves nor in LD with any causal sites—that is, at precisely those SNPs where otherwise an excess of false positives might occur.

We have focused on the case of confounding by parent genotype depicted in Figure 1 because of recently reported evidence for its occurrence (Kong et al., 2018; Lee et al., in press). In the Supplementary Note, we show that in the crucially distinct case of a parental trait affecting an offspring trait with which it is genetically *uncorrelated*, the intercept of LD Score regression does not reflect the degree to which such confounding inflates the GWAS statistics. A paradigm example of this case is a heritable nurturing trait such as intrauterine prenatal environment or milk yield affecting some distinct trait in the offspring, such as body size (Lynch & Walsh, 1998; Hadfield, 2012). To our knowledge, evidence concerning the existence of such “parental effects” in humans is scant. Studies of whether sharing a chorion tends to make monozygotic twins more similar have produced largely negative results for traits other birth weight (Marceau et al., 2016; van Beijsterveldt et al., 2016), and there have been several reports of null associations between parental behavior (maternal smoking, breastfeeding) and various behavioral outcomes (Der, Batty, & Deary, 2006; Lundberg et al., 2010; Skoglund, Chen, D’Onofrio, Lichtenstein, & Larsson, 2014). In the case of our motivating example, years of education, we think it highly plausible that the parental traits affecting offspring years of education that are much the same as the personal traits affecting one’s own educational attainment (intelligence, conscientiousness, openness, and so on) have much stronger quantitative impact than intrauterine prenatal environment, breastfeeding, or other forms of parental expenditure envisaged in the quantitative-genetic literature. There is theoretical and perhaps even practical interest in detecting any such parental effects that do exist, however, even if they are quantitatively small enough to neglect for purposes of LD Score regression. This is one motivation for increasing the sizes of family cohorts and phenotyping them more extensively.

Another type of scenario worth further consideration is batch effects (i.e., variability in genotyping/imputation pipeline within the same study) leading to spurious differences in allele frequencies between subgroups with genuinely different phenotypic means. It is not inconceivable that such confounding might be related to LD Score; for example, SNPs with higher LD Scores might be genotyped/imputed more accurately because they tend to have higher MAF and more redundant tagging. The additional question that must be asked, in this scenario and in any others that might offer themselves for contemplation, is whether the extent of such confounding at SNPs that are null by virtue of tagging few other SNPs of any kind can be safely generalized to a SNP with a larger LD Score that is nevertheless null because none of the many SNPs tagged by it happen to be causal. If this generalization is accurate or overly conservative, then the intercept of LD Score regression will continue to be a robust means of bringing the Type 1 error rate closer to the desired level. In the specific case of batch effects, we believe that the intercept is robust because such effects are more likely to afflict SNPs with lower LD Scores and thus lead to the intercept providing an overly conservative correction of null SNPs with higher LD Scores. Regardless of our own belief, however, the above-posed question of generalization from low-LD to high-LD null SNPs is what must be answered.

These conclusions depend importantly on the linearity of the relationship between LD Scores and the GWAS chi-square statistics (product of *Z* statistics). In real-data applications of LD Score regression to date, the chi-square vs. LD Score scatterplots have always borne out approximate linearity, and they should continue to be inspected in future applications. When users follow the developers’ recommendations for weighting of the SNPs in the regression, those SNPs with smaller LD Scores will receive larger weights, which in the case of nonlinearity brings the intercept closer to the conditional expected chi-square statistic of null SNPs. One form of nonlinearity that cannot be rectified by weighting is any misestimation of the theoretical intercept as a result of the slope being a biased estimate of heritability, but in practice this will lead to a spurious increase of the intercept and a consequent conservative correction of the GWAS statistics.

Even in cases where LD Score regression estimates the heritability (genetic covariance) with substantial bias, the method is able to estimate the genetic correlation with reasonable accuracy. Our mathematical analysis and simulation results suggest that the estimate should be treated with caution if it is statistically significant but nevertheless small. Our derivation in the Supplementary Note shows that any bias in the case of a small true genetic correlation will be minimal if the correlation depends primarily on direct overlap of the causal sites affecting the two traits—and negligibly on SNPs in LD with more potential causal sites thereby being more likely to tag one site affecting trait 1 and a distinct site affecting trait 2, with the signs of the alleles coupled with the reference allele at the tagging SNP showing a consistency across the genome. The reason why any bias resulting from the failure of this condition might be small, not exceeding 0.10 in magnitude, is that any such genome-wide pattern seems quite implausible; for example, if it is to create a substantial nonzero estimate of the genetic correlation when the true value is in fact zero, it amounts to causal sites that affect the two traits consistently occurring in the same genes and regulatory elements, with the appropriate coupling of alleles, but never coinciding. Furthermore, one might argue that this scenario (whatever its biological plausibility) does not necessarily invalidate bivariate LD Score regression as an estimator of the genetic correlation when this target quantity is defined properly. We have adopted the definition 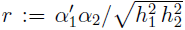 (see the Supplementary Note) in accordance with the Online Methods of Bulik-Sullivan et al. (2015b), but other authors have included contributions from LD and consistent coupling of allele signs to the definition of the genetic correlation (Lynch & Walsh, 1998).

A use of LD Score regression that we did not study in this work is the functional partition of heritability between different parts of the genome (Finucane et al., 2015). Simulation studies conducted by the authors suggest that this use is also quite robust, and this is probably the result of a similar cancellation of biases from numerator and denominator. A more recent work has introduced functional annotations describing many properties of SNPs related to per-SNP heritability, including MAF, local recombination rate, and extent of LD with neighbors (Gazal et al., 2017). Because these factors related to per-SNP heritability are thus effectively controlled, we might expect that the heritability estimate produced by stratified LD Score regression with these new annotations will be closer to the true heritability. Supplementary Table 8b of Gazal et al. (2017) does bear out a weak tendency for heritability estimates to increase in this manner. This same table, however, reveals a much stronger influence on heritability estimates; when stratified LD Score regression is applied to the summary statistics of a single large study rather than a meta-analysis of multiple studies, its heritability estimate becomes markedly higher and even approaches the estimate returned by the GREML method. Imperfect genetic correlations between studies thus seem to affect this output of GWAS as well (de Vlaming et al., 2017b). Applying stratified LD Score regression with the LD-related annotations to a large sample of a homogeneous population, analyzed with a uniform pipeline, appears to be promising strategy if the goal is to estimate heritability accurately. These conditions should also lead to an improved estimate of the intercept and a correction of the GWAS statistics that does not settle as much for being overly conservative (Loh et al., 2018).

In a field already marked by remarkable progress toward the goal of elucidating the causal relationship between its variables of interest without undue hindrance by confounding, LD Score regression adds a powerful new tool that allows many forms of confounding in a GWAS to be estimated and removed. In addition, it is a robust estimator of the genetic correlation, which is valuable in its own right because of its relevance to the causal nature of the phenotypic correlation (Duffy & Martin, 1994).

## Acknowledgements

This research was supported in part by the Intramural Research Program of the NIH, The National Institute of Diabetes and Digestive and Kidney Diseases (NIDDK).

